# HIV positive patients on antiretroviral therapy have an altered mucosal intestinal but not oral microbiome

**DOI:** 10.1101/2022.05.19.492600

**Authors:** Jingjing Meng, Junyi Tao, Yaa Abu, Daniel Andrew Sussman, Mohit Girotra, Sabita Roy

## Abstract

This study characterized compositional and functional shifts in the intestinal and oral microbiome in HIV-positive patients compared to uninfected individuals. 79 specimens were collected from 5 HIV-positive and 12 control subjects from five locations (colon brush, colon wash, TI brush, TI wash and saliva) during colonoscopy and at patient visits. Microbiome composition was characterized using 16S rRNA sequencing and microbiome function was predicted using bioinformatics tools (PICRUSt and Bugbase). Our analysis indicated that the β diversity of all intestinal samples (colon brush, colon wash, TI brush, TI wash) from patients with HIV was significantly different from patients without HIV, as measured by weighted UniFrac distances. Specifically, bacteria from genera *Prevotella, Fusobacterium, Eubacterium, Megasphaera, Mogibacterium* and *Mitsuokella* were more abundant in samples from HIV-positive patients. On the other hand, bacteria from genera *Ruminococus, Blautia*, and *Clostridium* were more abundant in samples from HIV-negative patients. Additionally, HIV-positive patients had higher abundances of biofilm-forming and pathogenic bacteria. Furthermore, pathways related to translation and nucleotide metabolism were elevated in HIV-positive patients whereas pathways related to lipid and carbohydrates metabolism and membrane transport were positively correlated with samples from HIV-negative patients. Our analyses further showed variations in microbiome composition in HIV-positive and negative patients by sampling site, with samples from colon wash, colon brush and TI wash significant between groups while samples from TI brush and saliva were not significant. Taken together, here we report altered intestinal microbiome composition and function in patients with HIV compared to uninfected patients, though we found no changes in the oral microbiome.

**Author summary:** Over 37 million people worldwide are living with HIV. Although the availability of antiretroviral therapy has significantly reduced the number of AIDS-related deaths, individuals living with HIV are at increased risk for opportunistic infections. We now know that HIV interacts with the trillions of bacteria, fungus, and viruses in the human body termed the microbiome. The advent of next generation sequencing has allowed us to characterize the composition of the microbiome and how HIV changes the microbiome composition to influence disease severity and progression. Previous studies have examined changes in the microbiome in HIV-positive and negative individuals. However, only a limited number of studies have compared variations in the oral and gastrointestinal microbiome with HIV infection. Furthermore, very few studies have looked at how the microbiome in HIV infection may vary by sampling site. Here, we detail how the oral and gastrointestinal microbiome changes with HIV infection and use 5 different sampling sites to gain a more comprehensive view of these changes by location. Our results show site-specific changes in the intestinal microbiome associated with HIV infection. Additionally, we show that while there are significant changes in the intestinal microbiome, there are no significant changes in the oral microbiome.

## Introduction

HIV (human immunodeficiency virus) infection is characterized by profound depletion of circulating and tissue-resident CD4+ T cells in gut associated lymphoid tissue and a chronic inflammatory state [1]. Although the overall survival of HIV patients has significantly improved since the introduction of antiretroviral therapy (ART), HIV-infected adults still have an increased risk of cardiovascular, liver, kidney, bone, and neurologic diseases [2] which are partially driven by microbial translocation and subsequent immune activation [3, 4].

In recent years, multiple groups have characterized the microbiome in the oral cavity or intestines in patients with HIV infection, though relatively few have examined both areas in the same patient. Although the oral cavity and the intestines are part of the gastrointestinal tract, they harbour distinct microbial communities, with the oral cavity dominated by *Firmicutes*, while the stool microbiota is mostly abundant in *Bacteroidetes* [5, 6]. These unique communities have been attributed to gastric acid in the stomach and bile acids in the duodenum [5, 7, 8]. Thus, while HIV is known to cause profound changes to the gastrointestinal system at large [9-11], recognizing site specific differences is key to fully appreciating the microbial landscape altered with HIV infection. Still, the majority of previous studies investigating intestinal microbial alterations with HIV infection only utilize one sampling site, with stool/stool swab [12, 13] samples being the most common, followed by rectal sponges [14] and rectosigmoid biopsy [15]. Similarly, in the oral cavity, the most common sampling sites utilized are saliva [16-19] or oral washes [20-22], with some studies using plaque samples [23, 24] or biofilm [25], but relatively few using multiple areas. Despite our understanding of site-specific microbial communities along the GI tract, to date, only one study has investigated the intestinal microbiome at different sites, including the terminal ileum, right colon, left colon, and feces [26], though the oral microbiome was not evaluated in this study.

Characterization of the intestinal microbiome in particular sheds light on the importance of sampling methods in identifying distinct microbial communities in many disease states. Gastrointestinal tract commensal bacteria consist of contents within the transient luminal compartment and the mucosal adherent compartment [27]. While most studies investigating the intestinal microbiota in humans have often used fecal samples because they are easily collected, the fecal microbiota is substantially variable between individuals and is often influenced by food/ingested materials, which limits our ability to identify specific disease associated microbes [27, 28]. On the other hand, the mucosa associated microbiota (MAM) is the more stable adherent compartment that adheres to mucosal surface of GI tract, though the main means of characterizing this compartment are through colonoscopic biopsies which is relatively invasive and limits their use [27, 28]. Notably, clinical studies of microbiome changes in HIV infection have shown differential bacterial microbiome phenotypes in intestinal biopsies, particularly in the terminal ileum, compared to fecal samples from the same individuals [26, 29]. These mucosal biopsies also have permitted examination of microbes that are most closely associated with the immune system [30]. Additionally, during colonoscopic biopsies, flushing of the mucosal surface with sterile water allows for the mucosal–luminal interface (MLI) to be sampled by washing off and collecting the loose mucus layer on the surface of the intestinal wall [28]. However, few studies have compared MLI sampling to biopsies, stool, or saliva.

In this study, we aim to characterize compositional and functional shifts in the mucosal intestinal microbiome and the oral microbiome in HIV-positive patients on ART compared to uninfected individuals using 16S rRNA sequencing. Here, we use multiple sampling sites, including brush samples and washes (colon brush, colon wash, terminal ileum brush, terminal ileum wash) during colonoscopic procedures as well as saliva samples at patient visits to gain a comprehensive view of intestinal and oral microbiome changes in the context of HIV infection.

## Results

### Patient selection

A total of 17 patients were enrolled for this study. 5 patients were diagnosed with HIV and 12 patients were HIV-negative. All HIV patients were on anti-retroviral therapy. Patient characteristics are summarized in Table 1 and detailed clinical characteristics for each subject can be found in Table S1. Notably, there was no significant difference in the mean age of patients between groups, though the female/male ratio and ethnicity in each group significantly differed.

**Table 1.** Patient characteristics.

|  | HIV-negative | HIV-positive |
| --- | --- | --- |
| **Number of subjects (n)** | 12 | 5 |
| **Mean age (years)** | 55 | 51 |
| **Female/male ratio** | 8/4 | 0/5 |
| **Ethnicity:** |  |  |
| **Hispanic (n)** | 8 | 4 |
| **Non-Hispanic (n)** | 4 | 1 |

Samples collected (Fig 1) in this study were obtained from terminal ileum (TI) wash, TI brush, colon wash, and colon brush of patients undergoing colonoscopy for comprehensive examination of mucosa associated microbiota (brush samples) as well as the mucosal–luminal interface (wash samples). Saliva samples were obtained directly from patients spitting into sterile containers to concomitantly survey the oral microbiome from the same patient.

**Fig 1.**
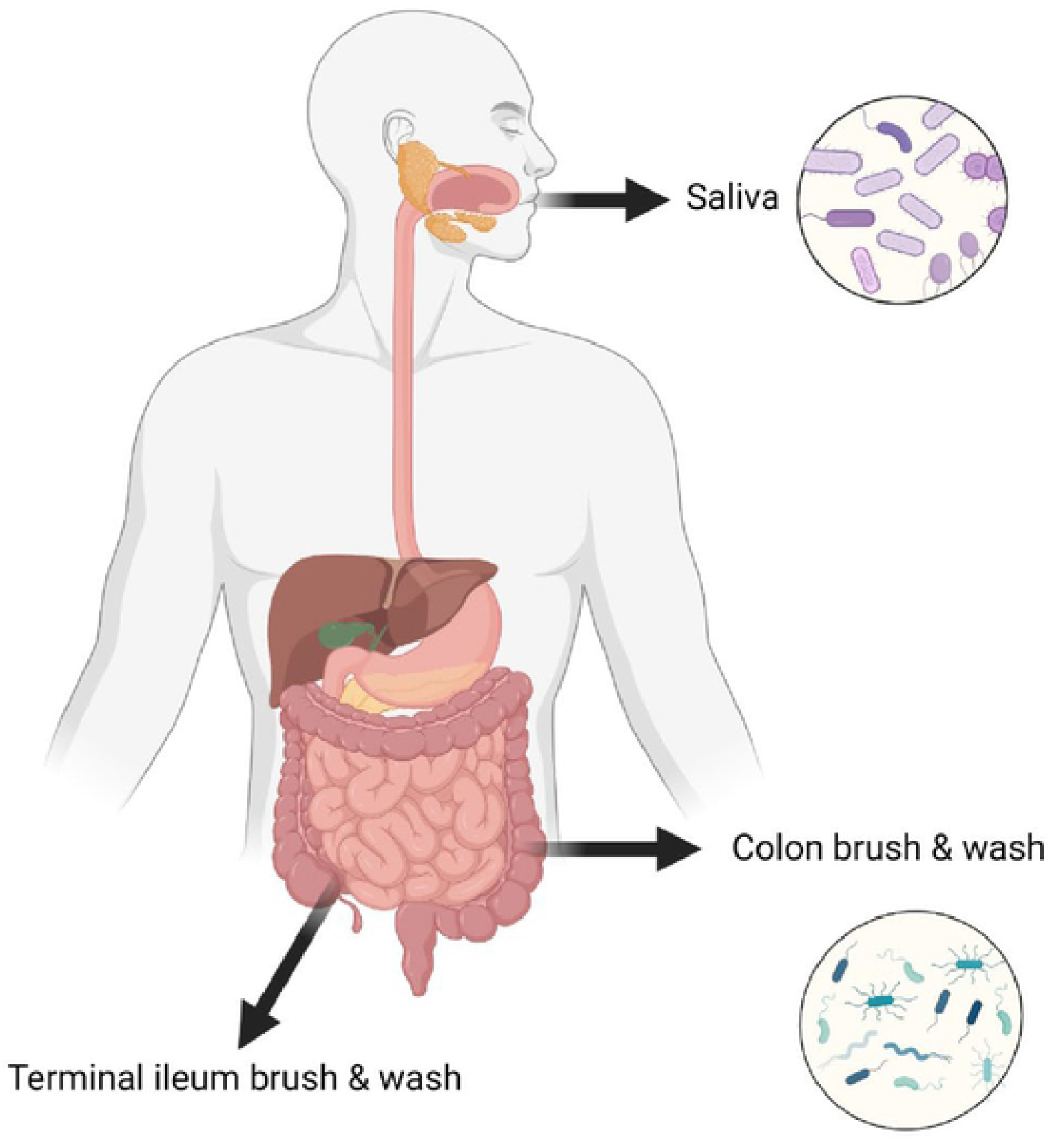
The schematics of all the sampling sites from human subjects.

### Patients with HIV have altered intestinal microbiome diversity and composition

On average, we obtained 178,608 sequence reads per intestinal sample and 176,926 sequence reads per saliva sample (S2 Table). Among all intestinal samples, 4310 unique ASVs were identified (S3 Table). Overall, our results show that HIV significantly alters the diversity of the intestinal microbiome. β diversity was assessed by weighted UniFrac distances and visualized with principal-coordinate analysis (PCoA) plots. Analysis of β diversity showed that all intestinal samples (colon brush, colon wash, TI brush, TI wash) from patients with HIV were significantly clustered apart from intestinal samples from patients without HIV (p < 0.05) (Fig 2A). Additionally, the α-diversity was measured by Faith’s phylogenetic diversity and Pielou’s Evenness. At sequencing depth of 80000, Faith’s phylogenic diversity was not significantly different between patients with or without HIV (p > 0.1) (Fig 2B). However, there was a tendency (p = 0.07) for intestinal samples from patients with HIV to exhibit higher evenness than those from patients without HIV (Fig 2B). Further, HIV infection also correlated with altered intestinal microbiome composition (S1 Fig). LefSe (Linear Discriminant Analysis (LDA) Effect Size) analysis was performed among intestinal samples to determine the bacterial taxa that were differentially enriched. Bacteria from genera *Prevotella, Fusobacterium, Eubacterium, Collinsella, Megasphaera, Mogibacterium* and *Mitsuokella* were more abundant in samples from HIV-positive patients. On the other hand, bacteria from genera *Ruminococus, Blautia*, and *Clostridium* are more abundant in samples from HIV-negative patients (Fig 2C).

**Fig 2.**
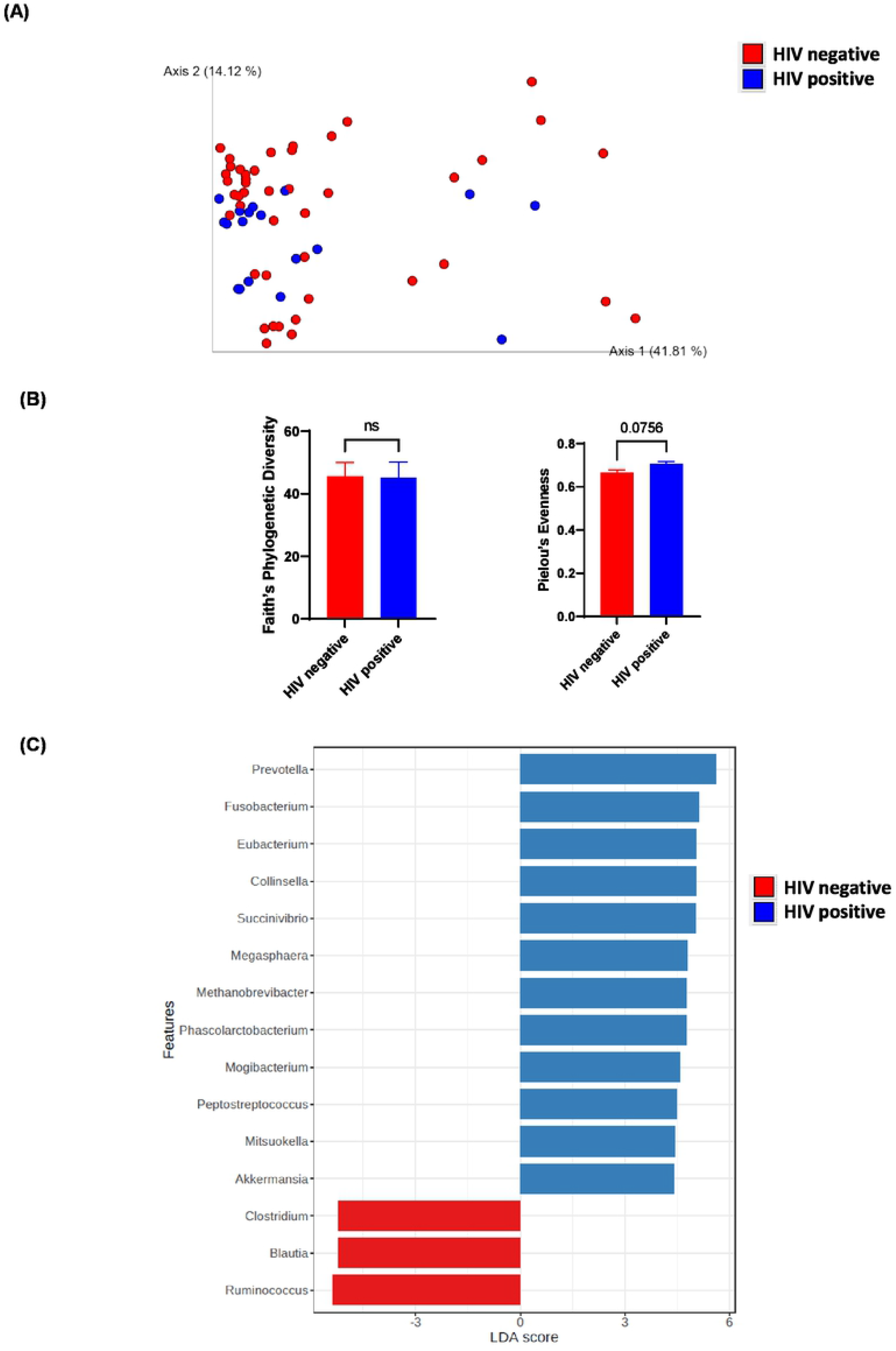
Diversity and composition analysis of the intestinal samples. Samples are grouped by HIV-negative (n=44) and positive (n=18). (A) Principal coordinates analysis (PCoA) plot of weighted UniFrac distances (metrics of β -diversity), p-value=0.012. (B) Faith Phylogenic Diversity and Pielou’s Evenness (metrics of α-diversity) at sequencing depth 80000. “ns” represents not significant, Error bars represent SEM. (C) LefSe (Linear Discriminant Analysis Effect Size) analysis of top discriminative bacteria genera between intestinal samples from HIV-positive and negative patients.

### Patients with HIV have altered intestinal microbiome function

BugBase algorithm was used to predict high-level phenotypes present in intestinal microbiome samples using 16S amplicon data. The BugBase phenotype predicted the abundances of gram-positive, gram-negative, biofilm-forming, and potentially pathogenic bacteria. Intestinal samples from HIV-positive patients had higher abundance of both biofilm-forming and pathogenic bacteria (p<0.01) (Fig 3A, B). Additionally, HIV-positive samples had higher percentages of gram-negative bacteria (p<0.01) and lower percentages of gram-positive bacteria (p<0.01) (Fig 3C, D).

**Fig 3.**
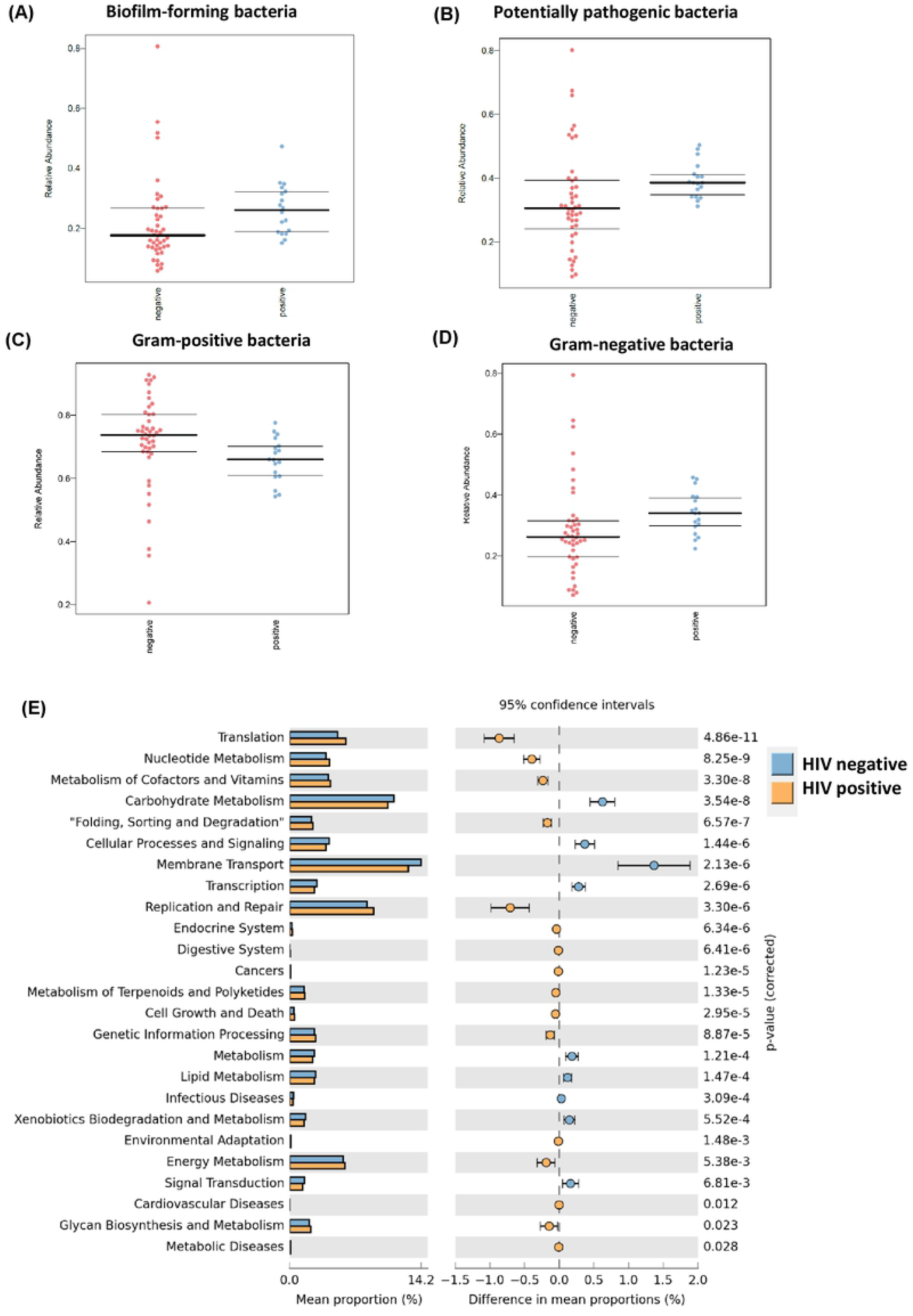
Predictive functional analysis of the intestinal samples. (A) BugBase predicted relative abundance of biofilm forming bacteria. Samples grouped by HIV-negative (n=44) and positive (n=18), p-value < 0.01. (B) BugBase predicted relative abundance of potentially pathogenic bacteria. Samples grouped by HIV-negative (n=44) and positive (n=18), p-value < 0.01. (C) BugBase predicted relative abundance of Gram-positive bacteria. Samples grouped by HIV-negative (n=44) and positive (n=18), p-value < 0.01. (D) BugBase predicted relative abundance of Gram-negative bacteria. Samples grouped by HIV-negative (n=44) and positive (n=18), p-value < 0.01. (E) The KEGG pathway of intestinal microbiota was predicted using PICRUSt (Phylogenetic Investigation of Communities by Reconstruction of Unobserved States). Data are presented in a barplot with 95% confidence intervals and p-values between intestinal samples from HIV-positive and negative patients.

The microbial metagenome was predicted with PICRUSt algorithm and functions were categorized with KEGG pathways to further elucidate the specific changes in microbial pathways. STAMP was used for identifying pathways that were differentially abundant between HIV-positive and negative patients. In total, 41 KEGG level-2 pathways were predicted among all intestinal samples (S4 Table). Pathways related to translation, nucleotide metabolism, cofactors and vitamin metabolism, and replication and repair were positively correlated with samples from HIV-positive patients (Fig 3E). On the other hand, pathways related to lipid and carbohydrates metabolism, membrane transport, signalling transduction, and cellular processes and signalling were positively correlated with samples from HIV-negative patients (Fig 3E).

### Differences in HIV-associated intestinal microbiome diversity and composition by sampling site

Intestinal samples were collected from 4 different sampling sites. Following, we then compared samples from HIV-positive and HIV-negative patients at each sampling site (colon wash, colon brush, TI wash, and TI brush). In colon wash samples, the β diversity, measured using unweighted UniFrac distances, between HIV-positive and HIV-negative samples was significantly different (p < 0.05) (Fig 4A). Additionally, in colon wash, HIV-positive samples had higher richness (P<0.01) than HIV-negative samples when the α-diversity was measured by Faith’s phylogenetic diversity (Fig 4B). However, there was no difference in sample evenness between HIV-positive and negative colon wash samples (Fig 4B). Furthermoe, LefSe analysis showed that bacteria from genera *Succinivibrio* and *Aggregatibacter* was more abundant in HIV-positive colon wash samples (Fig 4C).

**Fig 4.**
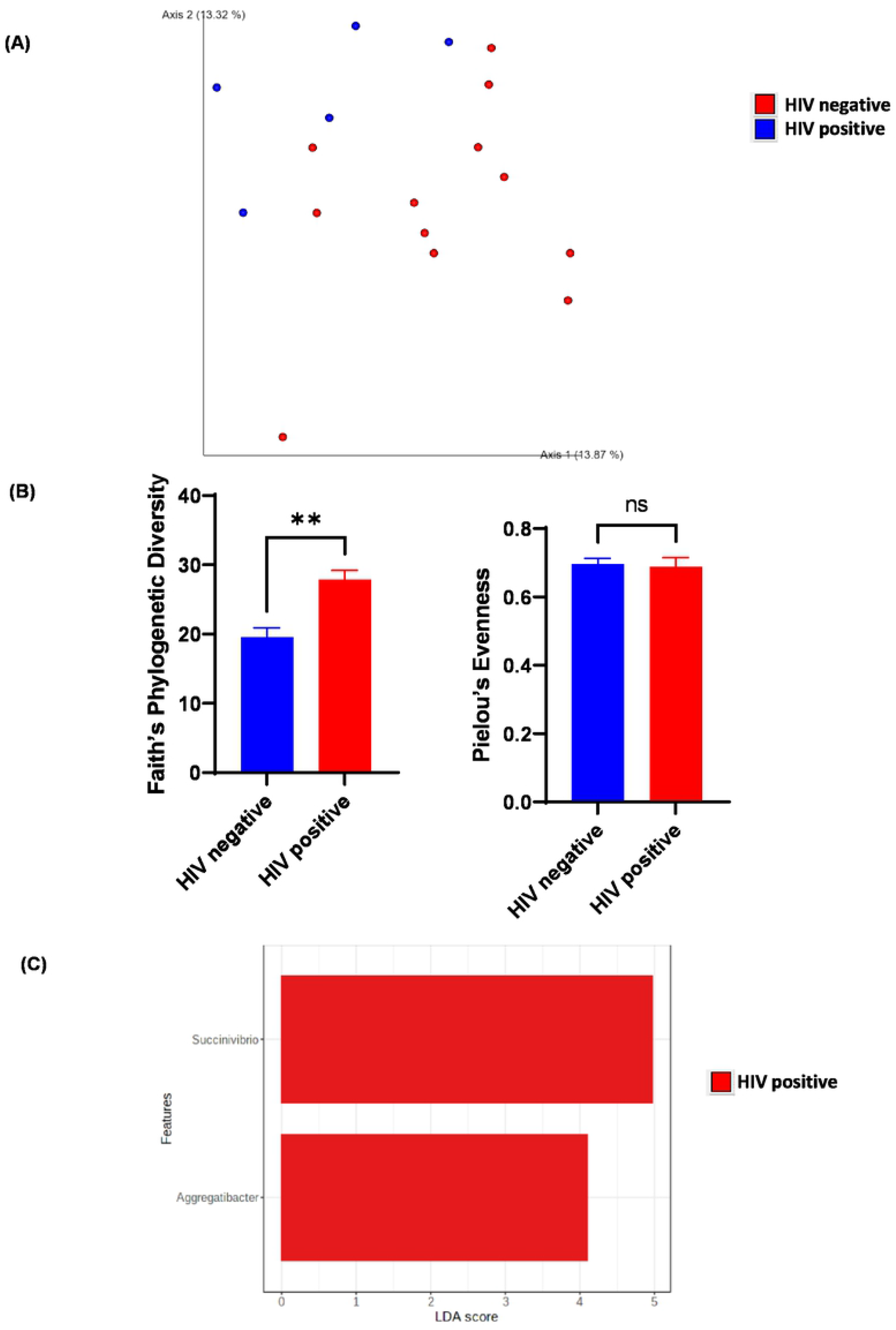
Diversity and composition analysis of the colon wash samples. (A) Principal coordinates analysis (PCoA) plot of unweighted UniFrac distances (metrics of β - diversity). Samples grouped by HIV-negative (n=12) and positive (n=5), p-value=0.013. (B) Faith Phylogenic Diversity and Pielou’s Evenness (metrics of α-diversity) at sequencing depth 80000. Samples grouped by HIV-negative (n=12) and positive (n=5). “ns” represents not significant, “**” represents P< 0.001. Error bars represent SEM. (C) LefSe (Linear Discriminant Analysis Effect Size) analysis of top discriminative bacteria genera between intestinal samples from HIV-positive and negative patients.

In colon brush samples, HIV-positive and HIV-negative samples showed a tendency to cluster differently (p =0.073) when β diversity was measured using weighted UniFrac distances (Fig 5A). Additionally, there was no difference in sample richness or evenness between colon brush samples (Fig 5B). Notably, bacteria from *Megasphaera* and *Slackia* were enriched in HIV-positive colon brush samples (Fig 5C).

**Fig 5.**
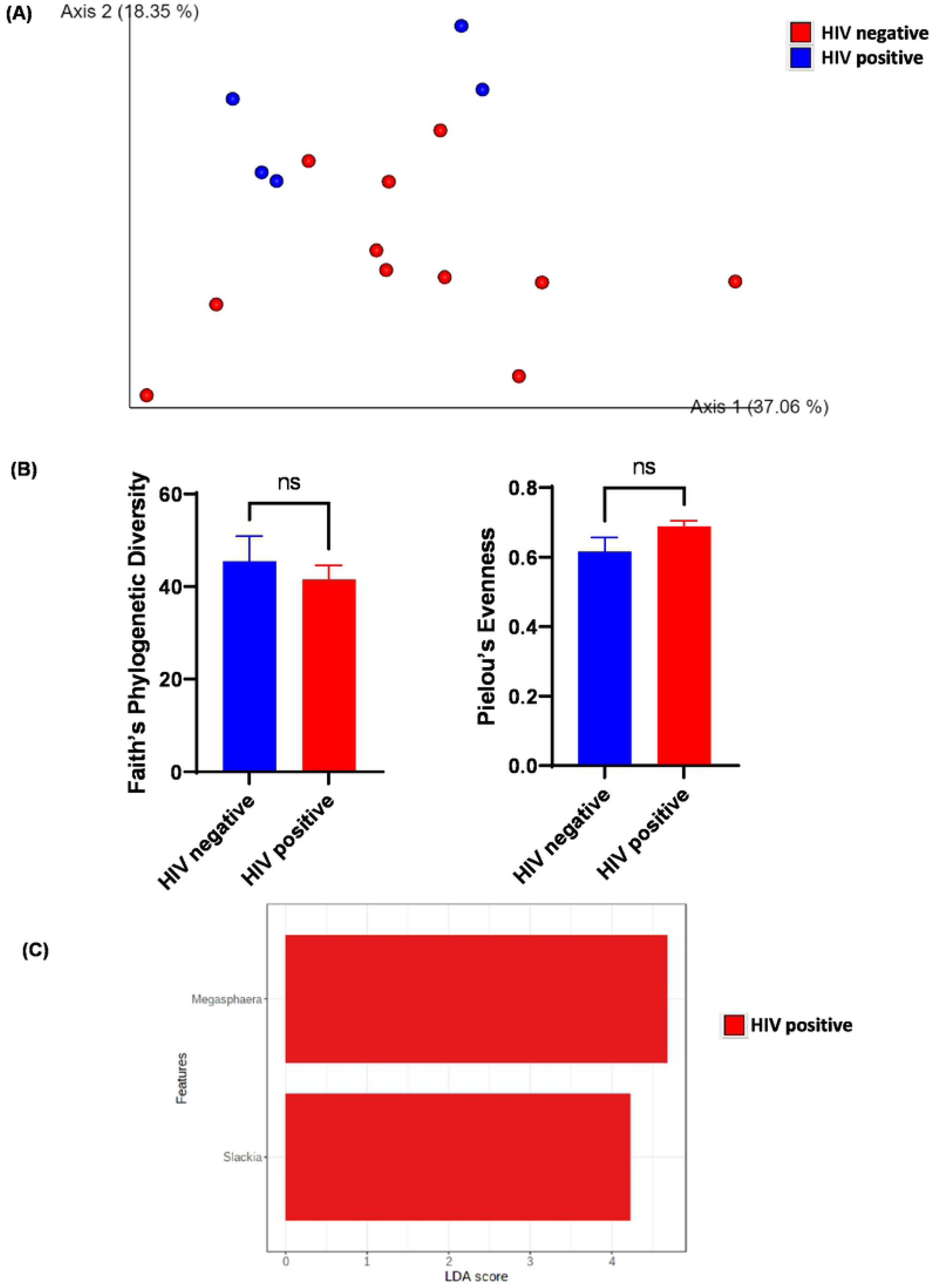
Diversity and composition analysis of the colon brush samples. (A) Principal coordinates analysis (PCoA) plot of weighted UniFrac distances (metrics of β - diversity). Samples grouped by HIV-negative (n=12) and positive (n=5), p-value=0.073. (B) Faith Phylogenic Diversity and Pielou’s Evenness (metrics of α-diversity) at sequencing depth 80000. Samples grouped by HIV-negative (n=12) and positive (n=5). “ns” represents not significant, bars represent SEM. (C) LefSe (Linear Discriminant Analysis Effect Size) analysis of top discriminative bacteria genera between intestinal samples from HIV-positive and negative patients.

In TI wash samples, HIV-positive and HIV-negative samples were significantly different (p < 0.05) as assessed by unweighted UniFrac distances (Fig 6A). However, there was no difference in sample richness or evenness between TI wash samples (Fig 6B). Notably, bacteria from *Prevotella, Mogibacterium, Mitsuokella* and *Aggregatibacter* were enriched in HIV-positive TI wash samples (Fig 6C).

**Fig 6.**
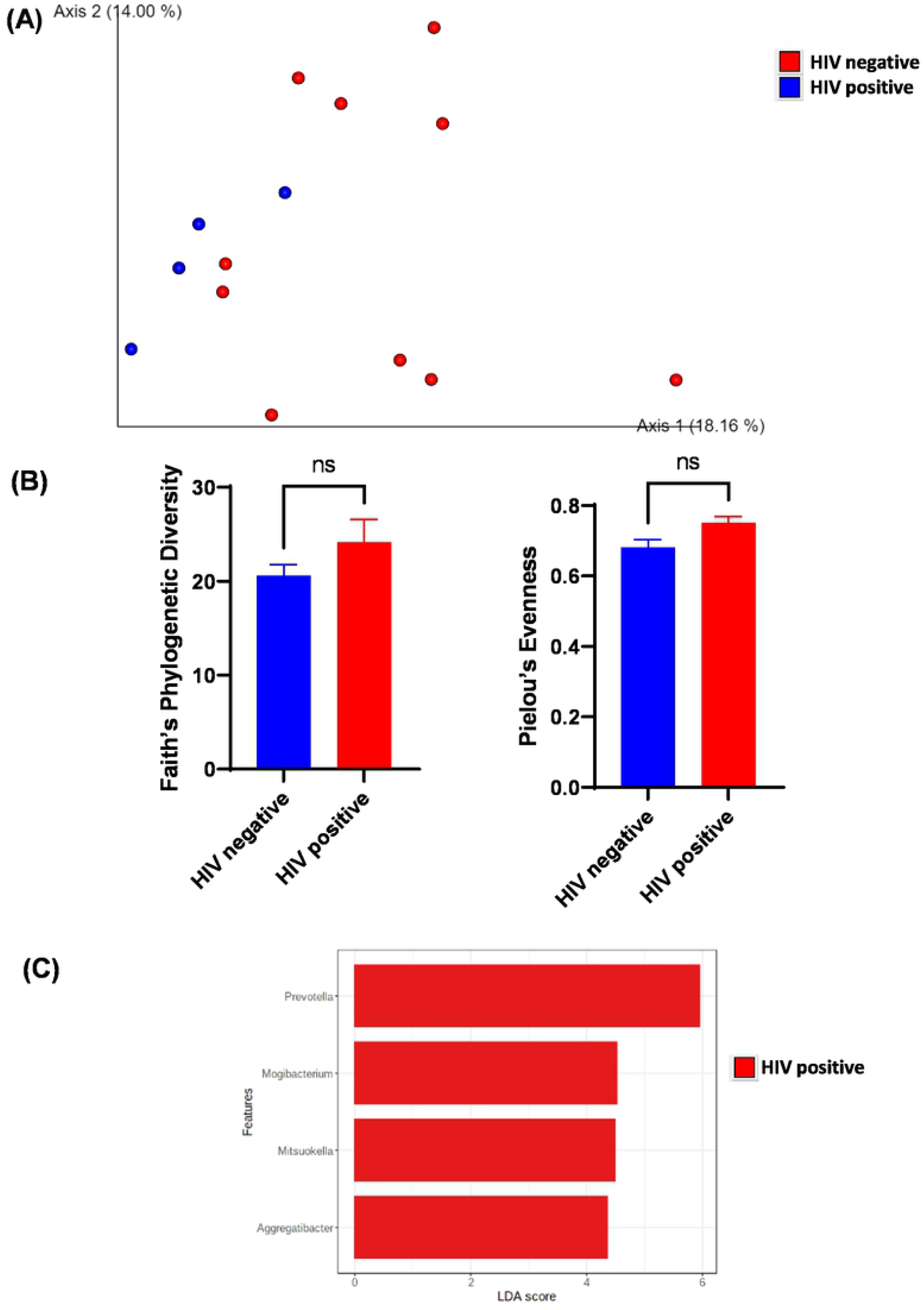
Diversity and composition analysis of the TI wash samples. (A) Principal coordinates analysis (PCoA) plot of unweighted UniFrac distances (metrics of β - diversity). Samples grouped by HIV-negative (n=12) and positive (n=5), p-value=0.02. (B) Faith Phylogenic Diversity and Pielou’s Evenness (metrics of α-diversity) at sequencing depth 80000. Samples grouped by HIV-negative (n=12) and positive (n=5). “ns” represents not significant.Error bars represent SEM. (C) LefSe (Linear Discriminant Analysis Effect Size) analysis of top discriminative bacteria genera between intestinal samples from HIV-positive and negative patients.

In TI brush samples, HIV-positive and HIV-negative samples were not significantly different (p > 0.2) as assessed by weighted UniFrac distances (Fig 7A). Additionally, there was no significant difference in sample richness or evenness between TI wash samples between groups (Fig 7B). Moreover, LefSe analysis showed no bacteria taxa enriched in either group.

**Fig 7.**
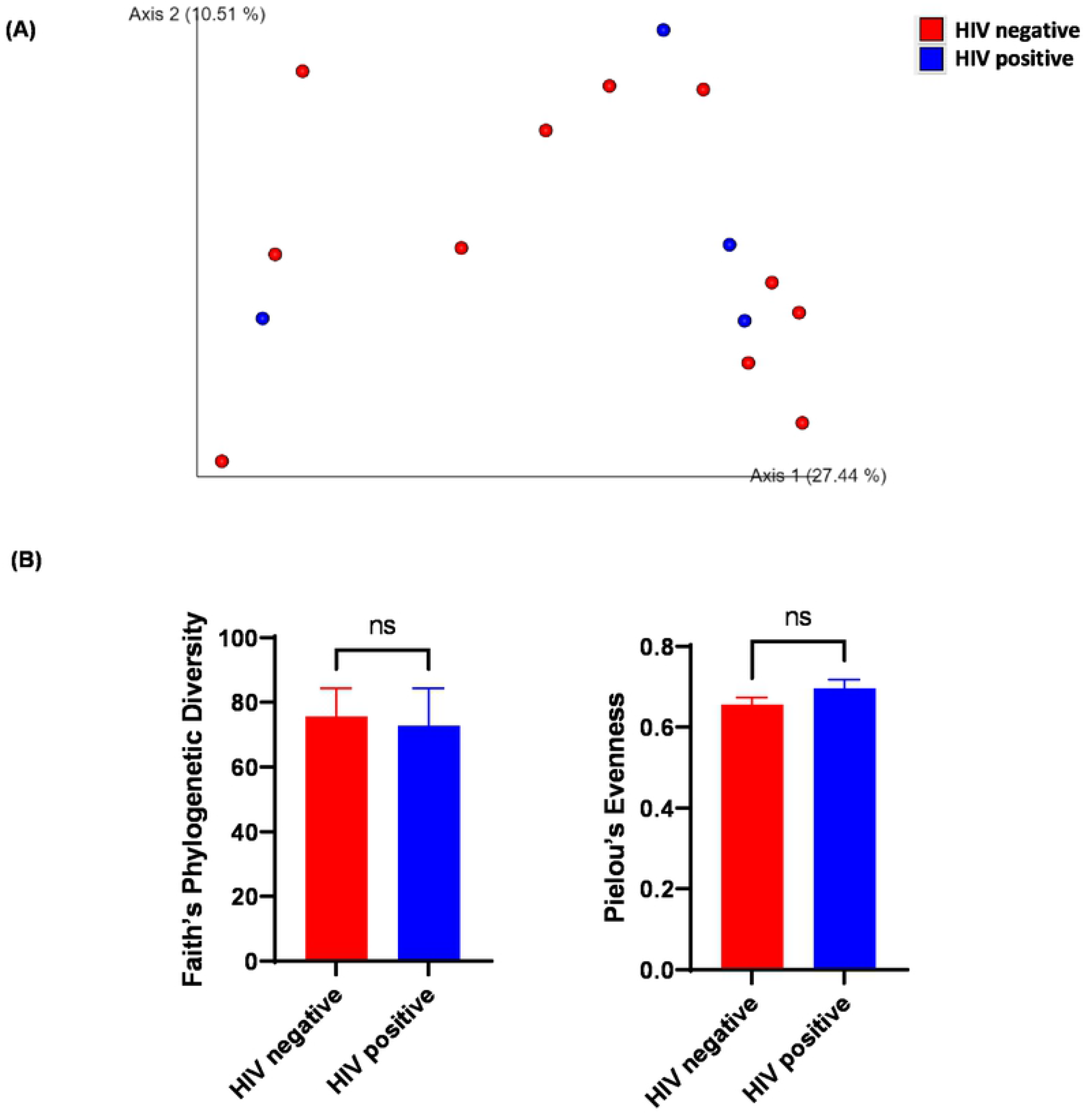
Diversity analysis of the TI brush samples. (A) Principal coordinates analysis (PCoA) plot of weighted UniFrac distances (metrics of β -diversity). Samples grouped by HIV-negative (n=11) and positive (n=4), p-value>0.2. (B) Faith Phylogenic Diversity and Pielou’s Evenness (metrics of α-diversity) at sequencing depth 80000. Samples grouped by HIV-negative (n=11) and positive (n=4), p -value >0.2. “ns” represents not significant.

To investigate the impact of sampling sites on the microbiome, we also analyzed the samples from HIV-negative patients alone. TI brush samples significantly clustered apart from the three other sampling locations (TI wash, colon wash, and colon brush) in the PCoA plot (q <0.05) (S3 Fig). Additionally, both TI and colon brush samples had higher Faith’s phylogeny diversity than TI and colon wash samples (q <0.05) (S3 Fig), indicating more unique bacteria taxa are present on the intestinal epithelial. Additionally, there was no significance between TI and colon wash samples assessed by Faith’s phylogeny diversity.

### The oral microbiome is not altered in patients with HIV

Among all oral samples, 1520 unique ASVs were identified (S3 Table). The composition of the oral microbiome was plotted at the phylum and genus level and grouped by HIV status (S2 Fig). Our results indicate that the oral microbiome is not altered with HIV infection. When β diversity was measured using weighted UniFrac distances and visualized with PCoA plots, salivary samples from patients with HIV did not cluster apart from salivary samples from patients without HIV (p >0.1) (Fig 8A). Furthermore, neither Faith’s phylogenetic diversity nor Pielou’s Evenness demonstrated differences between HIV-positive and HIV-negative saliva samples (p >0.1) (Fig 8B). Notably, the pCoA plot of HIV-negative samples also confirmed that the oral microbiome is significantly different from the intestinal microbiome (q<0.01) (S3A Fig).

**Fig 8.**
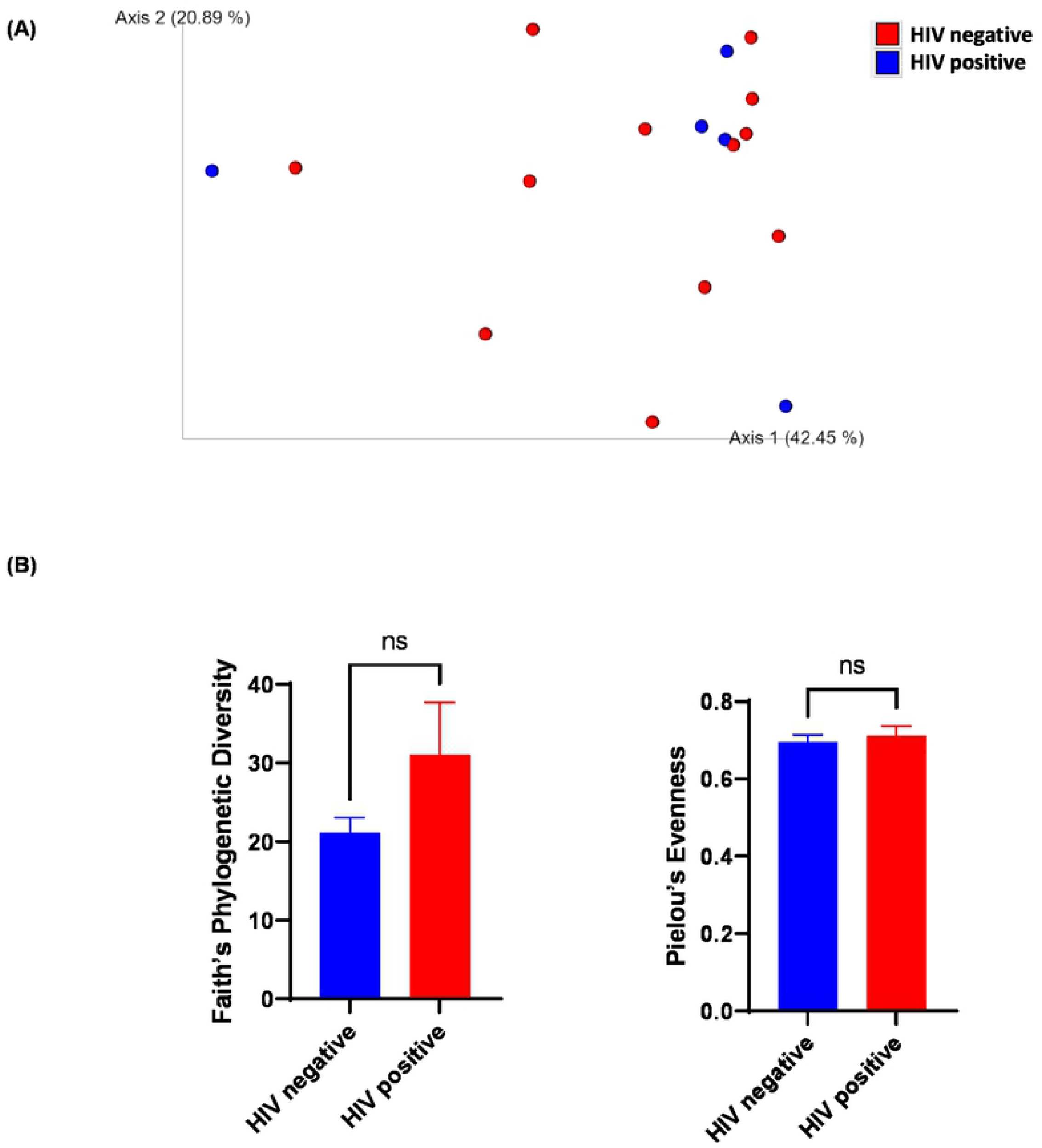
Diversity analysis of the saliva samples. (A). Principal coordinates analysis (PCoA) plot of weighted UniFrac distances (metrics of β -diversity). Samples grouped by HIV-negative (n=12) and positive (n=5), p-value>0.2. (B) Faith Phylogenic Diversity and Pielou’s Evenness (metrics of α-diversity) at sequencing depth 80000. Samples grouped by HIV-negative (n=12) and positive (n=5), p -value >0.2. “ns” represents not significant.

## Discussion

In this study, we characterized both the oral and intestinal microbiome using multiple sampling sites (colon brush, colon wash, TI brush, TI wash) in patients with HIV, which has not previously been described in samples from the same patients. Here, we report alterations in the composition and function of the intestinal microbiome in HIV-infected patients compared to uninfected individuals. Interestingly, we found no differences in the composition and function of the oral microbiome between groups. In our study, we also compared the intestinal microbiome by sampling site and demonstrate site-specific alterations in the microbiome, with the microbiome unaltered in TI brush samples, while other sites showed significant differences.

Our findings regarding the β diversity in intestinal samples of HIV patients were consistent with previous studies [9, 10, 31], which also demonstrate a shift in in the overall intestinal microbial community. In terms of α diversity, this study did not find a significant difference in sample richness or evenness; however, a tendency of increased evenness was observed. Consistently, Vujkovic-Cvijin et al. reported no changes in community richness or evenness [15] and Dillon et al. observed similar sample richness and evenness between uninfected and HIV infected subjects and a trend towards greater evenness in HIV infected individuals [29]. However, other studies have reported either an increase or decrease in alpha diversity in HIV patients [13, 14, 32, 33]. Additionally, Lozupone et al. reported that untreated HIV patients had higher alpha diversity [34]. The discrepancies in results are likely due to variations in diet associated with different regions and ethnicities, relatively small sample sizes in each study, and lack of proper control subjects.

We also identified differentially enriched bacteria with HIV infection and by sampling site. Several studies [29, 34, 35] have reported enrichment of *Prevotella* in HIV positive subjects, which is in agreement with our study that *Prevotella* had the highest LDA score in LefSe analysis. Interestingly, one study reported the *Prevotella* was significantly decreased after ART, suggesting its involvement during HIV inflammation [13]. Furthermore, Noguera-Julian et al. observed that the balance between *Prevotella* and *Bacteriodes* may be correlated with sexual preferences rather than HIV per se [36]. Specifically, they found that men who have sex with men (MSM) had increased *Prevotella*, whereas most non-MSM subjects were enriched in *Bacteroides*, regardless of HIV-1 status. Notably, many of our identified bacteria enriched in HIV patients were consistent with previous studies. Mchardy et al. reported *Fusobacteria* and *Peptostreptococcus* among the bacteria that were significantly enriched in HIV-positive patients[14]. Lozupone et al. also reported that *Peptococcus, Mitsuokella jalaludinii* and *Megasphaera elsdenii* increased in relative abundance in patients with HIV infection [34, 37]. Mutlu et al. observed an increase in *Mogibacterium* and unclassified *Fusobacteriaceae* associated with HIV [26]. However, there was discordance in our studies and others regarding *Eubacterium*, with two studies reporting that *Eubacterium* were depleted in HIV associated mucosal samples [14, 26]. Discrepancies could be attributed method for sample collection/reservation, diet effect associated with different regions and ethnicities. Still, the bacteria genera we identified that were more enriched in non-HIV patients are consistent with previous literature. For example, McHardy et al. and Mutlu et al. reported that *Ruminococcus* was depleted in HIV-infected subjects[14, 26], which is in accordance with our finding that *Ruminococus* is more abundant in HIV-negative patients. Consistently, others have reported genus *Blautia* were decreased in HIV patients in accordance with our own observations [26, 29].

Previous studies also examined functional shifts in the intestinal metagenome in HIV infected individuals. McHardy et al. reported that the imputed microbial metagenome from HIV patients without ART treatment were depleted of amino acids metabolism, CoA biosynthesis and fructose/mannose metabolism; and were enriched for glutathione metabolism, selenocompound metabolism, folate biosynthesis, and siderophore biosynthesis[14]. Additionally, Vázquez-Castellanos et al did metagenome sequencing on the intestinal microbiota and found enrichment of genes involved in various pathogenic processes, lipopolysaccharide biosynthesis, bacterial translocation, and other inflammatory pathways in HIV-positive individuals. Furthermore, genes involved in amino acid metabolism and energy processes were depleted in HIV-positive individuals[1]. Our own observations of altered microbiome functions largely agreed with previous studies. BugBase and PICRUSt functional algorithms predicted that potentially pathogenic bacteria and Gram-negative bacteria were enriched in HIV-positive patients. Similarly, our functional profiling predicted depletion of carbohydrates and lipids metabolism in HIV-positive individuals, which are both related to overall energy metabolism. However, we did not observe significant changes in amino acids metabolism in our study, which could be partially attributed to the diet of the study subjects.

To date, very few studies have investigated changes in the intestinal microbiome using multiple samplings sites. One study concluded that fecal aspirates and stool samples generally represent the same pattern of bacteria from *Bacteroidetes* in mucosa-adherent bacteria, but HIV-1-associated changes in *Proteobacteria* and *Firmicutes* were mucosa-specific [29]. Furthermore, Yang et al found that the difference between HIV positive and negative groups was not significant when all four body sites (mouth, esophagus, stomach and duodenum) were included, and the difference became significant when the proximal intestinal (esophagus, stomach and duodenum) was analyzed. In site-specific analyses, separation between HIV patients and controls was only significant in the duodenum but not significant in each of the three other body sites examined [38]. Mutlu et al. reported that all sample types (terminal ileum, right colon, left colon, feces) had less OTUs in the HIV group compared to healthy controls. Additionally, they observed separation of samples for each sample type when β diversity were measured by UniFrac metric. The dispersion of samples was visually more apparent for the terminal ileum and right colon samples, and less overlapping in the left colon and fecal samples [26]. Our study also reflects sampling-site specific variations in the intestinal microbiome. Samples from colon wash, colon brush and TI wash were significantly different in microbial composition between HIV infected and uninfected subjects, while samples from TI brush and saliva were not significant. Moreover, we demonstrated that TI brush samples were very different from the three other sampling locations (TI wash, colon wash, and colon brush) in the β diversity plot (S3 Fig). Our study also highlights differences in alpha diversity in sampling site (lumen wash vs. lumen brush), as both TI and colon brush samples had higher Faith’s phylogeny diversity than TI and colon wash samples (S3 Fig), indicating more unique bacteria taxa are present on the intestinal epithelial. Overall, our results suggest that HIV infection affected the microbiome in a site-specific manner.

Previous studies have also profiled the oral microbiome with HIV infection. However, there have been inconsistent reports on the effect on alpha diversity. For instance, one group reported that the oral microbiota in HIV infected patients had higher alpha diversity as well as higher bacterial loads[19]. However, others reported a lower oral microbiome richness in HIV-infected individuals [21, 39]. The increase in alpha diversity could be the result of an increase in pathogen colonization, and the decrease could be the result of a few pathogens that dominated the oral environment. Further, a range of secretory antimicrobial peptides could also play a significant role in the balance[40]. That being said, while several studies have demonstrated that the oral microbiome composition changes during HIV infection[16, 39], other studies report no major change in the oral microbiome or attribute changes to co-morbid periodontal disease [21, 23, 41]. In the current study, we did not observe any significant change in oral microbiome richness, evenness, or composition.

Collectively, our study highlights site-specific alterations in the microbiome, and support the possibility of targeting certain regions in the gastrointestinal tract to mitigate dysbiosis in HIV-positive patients. Additionally, the use of brush and wash samples allowed us to determine changes in adherent or loosely associated mucosa microbiota. Broadly, our findings add to the general knowledge to aid the development of precise-location and microbial targeted interventions. This study also had several limitations, which must be noted. First, this study had a small sample size (HIV+ = 5 individuals; HIV- = 12 individuals), reflecting difficulty in recruiting subjects for biopsy and/or poor adherence with patients follow up visits for saliva samples. A larger sampling size would have permitted more differentially abundant bacteria taxa to be detected. Next, this study did not correlate microbiome findings with immune measurements. Importantly, the correlation of CD4+ T cells has often been associated with specific bacteria taxa, and these associations were lacking in this study.

As this study collected samples from biopsy rather than stool, microbiome analyses presented here are less biased by the substantial variations caused by food/ingested materials in human subjects. Still, it must be noted that some studies investigating the intestinal and oral microbiome with HIV have been inconsistent in their metrics, which has largely been attributed to diet. For instance, some studies have reported patients with HIV have lower alpha diversity [13, 14, 26] in intestinal samples; however, others claim HIV patients have higher [34], or no change in intestinal alpha diversity [15, 29]. Similarly, there have been differences in enriched bacterial taxa across studies in intestinal samples in HIV patients. Some groups reported depletion of *Bacteroides* and enrichment of *Prevotella* in HIV [26, 34]. Others reported depletion of *Clostridiales* in untreated HIV patients [14]. Furthermore, increased *Proteobacteria* and decreased *Firmicutes* in HIV have also been reported [38]. Additionally, there have been inconsistencies in the reporting of the oral microbiome between HIV infected patients and healthy controls [42]. Jimenez-Hernandez *et al*. reported that the salivary alpha diversity in HIV-infected individuals were significantly higher than those in HIV-uninfected samples [19]. Others revealed that the oral alpha diversity in patients with HIV was significantly lower than uninfected individuals [16, 21, 39]. Thus, we consider our approach using biopsies to profile the mucosal microbiota at specific sites within the gastrointestinal tract and saliva samples to characterize the oral microbiome more advantageous compared to these aforementioned studies.

In conclusion, in the current study, we show altered intestinal microbiome composition and function in patients with HIV, with no significant differences in the oral microbiome between HIV infected and uninfected patients. Here, we also characterized changes in the intestinal microbiome by site of sampling and found that samples from colon wash, colon brush and TI wash were significant between groups while samples from TI brush and saliva were not significant. As the role of the microbiota is becoming increasingly clear in HIV infection, our study, which profiles the oral microbiota, strongly adherent mucosal communities (brush), and loosely mucosa-associated microbiota (wash) help put into context site-specific changes with HIV infection.

### Patients and Methods

#### Study subjects and sample collection

This work was conducted with approval from University of Miami’s Institutional Review Board (20160338). Inclusion criteria were individuals between 18-65 years old who consented to this study and were scheduled to undergo endoscopy and/or colonoscopy or sigmoidoscopy at either the University of Miami Hospital or the University of Miami Hospital & Clinics/Sylvester Comprehensive Cancer Center. HIV-positive patients were defined as having a previous diagnosis of HIV per electronic medical record. Patients provided >5mL saliva during hospital visits, using the Omnigene Oral kit (DNA genotek, OM-501). TI brush and colon brush were collected during colonoscopy or sigmoidoscopy with a sheathed cytology brush (ConMed, 000110). All samples were stored at -80°C until processing.

### DNA extraction and 16S rRNA gene sequencing

For intestinal samples (colon brush, colon wash, TI brush, TI wash), DNA was isolated using DNeasy PowerSoil Pro Kit (Qiagen, 47016). For saliva samples, DNA was isolated using QIAamp DNA Blood Mini Kit (Qiagen, 51104). During DNA extraction, two extraction controls were included to remove potential contamination from kit reagents. Sequencing was performed by the University of Minnesota Genomics Center. The hypervariable regions V4 region of 16S rRNA gene was PCR amplified using the forward primer 515F (GTGCCAFCMGCCGCGGTAA), reverse primer 806R (GGACTACHVGGGTWTCTAAT), Illumina adaptors, and molecular barcodes to produce 427 base pair (bp) amplicons. Amplicons were sequenced with the Illumina MiSeq v.3 platform, generating 300-bp paired-end reads. The extraction controls could not be PCR amplified and were therefore excluded from the sequencing process.

### Bioinformatics analysis

Demultiplexed sequence reads were clustered into amplicon sequence variants (ASVs) with the DADA2 package (version 1.21.0) [43] implemented in R (version 4.0.3) and RStudio (version 1.1.463). The steps of the DADA2 pipeline include error filtering, trimming, learning of error rates, denoising, merging of paired reads, and removal of chimeras. On average, 178,608 sequence reads per intestinal sample and 176,926 sequence reads per saliva sample were kept after error filtering and other steps (S2 Table). During trimming, the forward and reverse reads were truncated at positions 230 and 180 to remove low-quality tails. The ASV table generated by DADA2 was imported into the QIIME2 pipeline[44] for diversity analyses and taxonomic assignment. Diversity analyses were performed by using the qiime diversity core-metrics-phylogenetic script with sampling depth of 80,000. Taxonomic assignment of ASVs was done to the genus level using a naive Bayesian classifier[45] implemented in QIIME2 with Greengenes reference database (13_8 99%)[46]. MicrobiomeAnalyst[47] was used for generating bar plots and LefSe (Linear discriminant analysis Effect Size)[48] plot. The threshold on the logarithmic LDA score for discriminative features was set to 2. PICRUSt[49] is a computational approach to predict the functional composition of a metagenome using 16S data with reference genomes from Greengenes[46] and IMG[50] databases. PICRUSt pathway prediction was implemented within galaxy app (https://huttenhower.sph.harvard.edu/galaxy/). KEGG orthologs [51] was used to predict metagenome. KEGG pathway was categorized to pathway hierarchy level 2. STAMP [52] was used for identifying pathways that were differentially abundant and for generating extended error bar plot. BugBase [53] is a microbiome analysis algorithm that predicts high-level phenotypes present in microbiome samples using 16S amplicon data. The BugBase phenotype predictions were implemented using the online web app (https://bugbase.cs.umn.edu/). The False discovery rate (q-value) threshold was set to 0.05.

### Statistical analysis

Mann-Whitney test or Kruskal-Wallis test was used to detect if α diversity differed across treatments. Permutational multivariate analysis of variance (PERMANOVA) was used to detect if β diversity differed across treatments. Benjamini-Hochberg method was used for controlling false discovery rate (q-value). P < 0.05 was considered to be statistically significant.

## Data availability

Sequence data are available at the Biostudies database [54] (https://www.ebi.ac.uk/biostudies/) under accession number S-BSST836.

## Ethics statement

The study protocol was approved by The University of Miami Institutional Review Boards (20160338). Written and informed consent was obtained from each patient before enrolment. All patients were enrolled at the University of Miami Hospital.

## Supporting Information

**S1 Fig.**
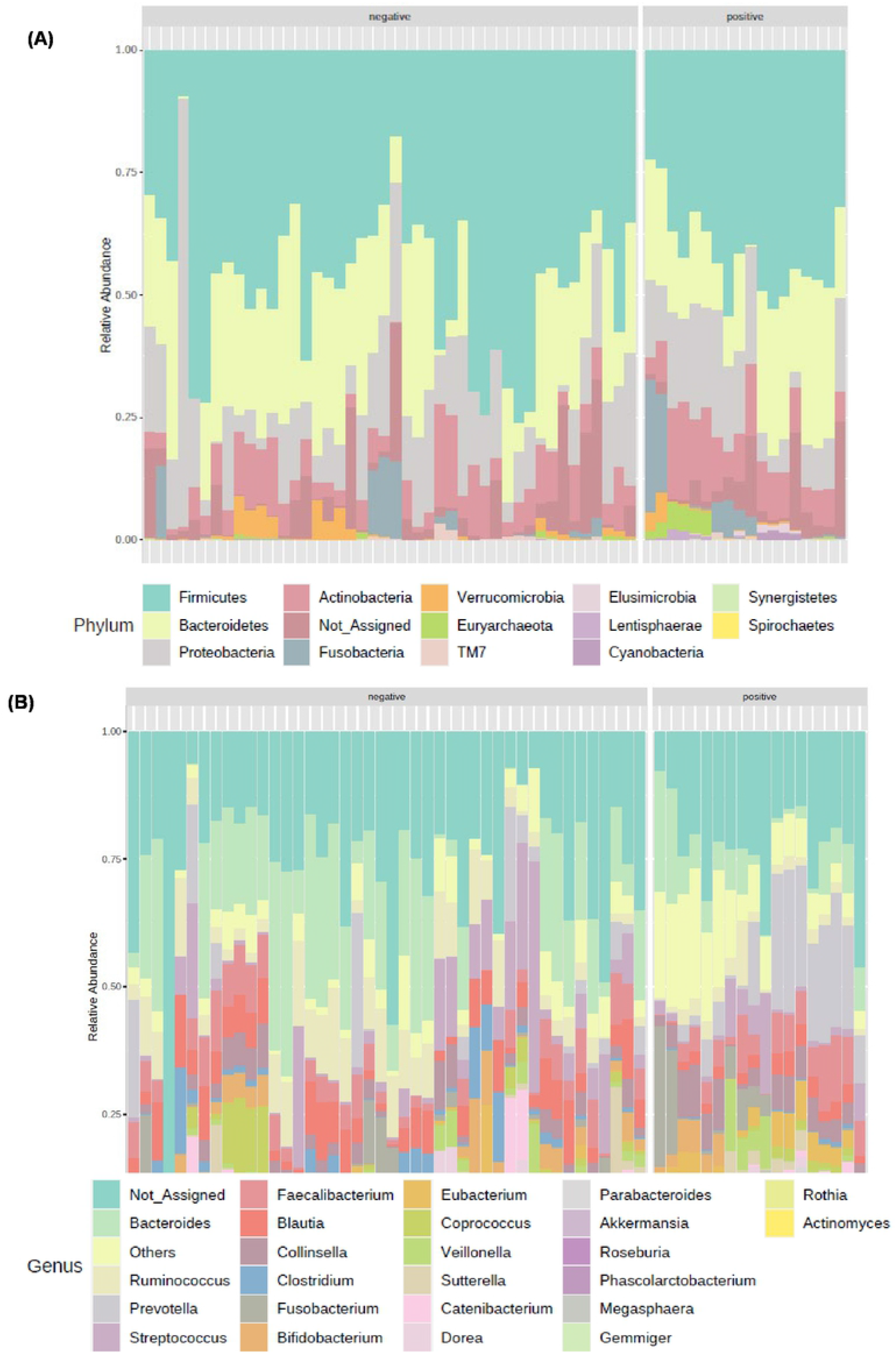
Bar plots of bacterial composition in the intestinal samples. Samples were grouped by HIV-negative (n=44) and positive (n=18). (A) at the phylum level. (B) at the genus level.

**S2 Fig.**
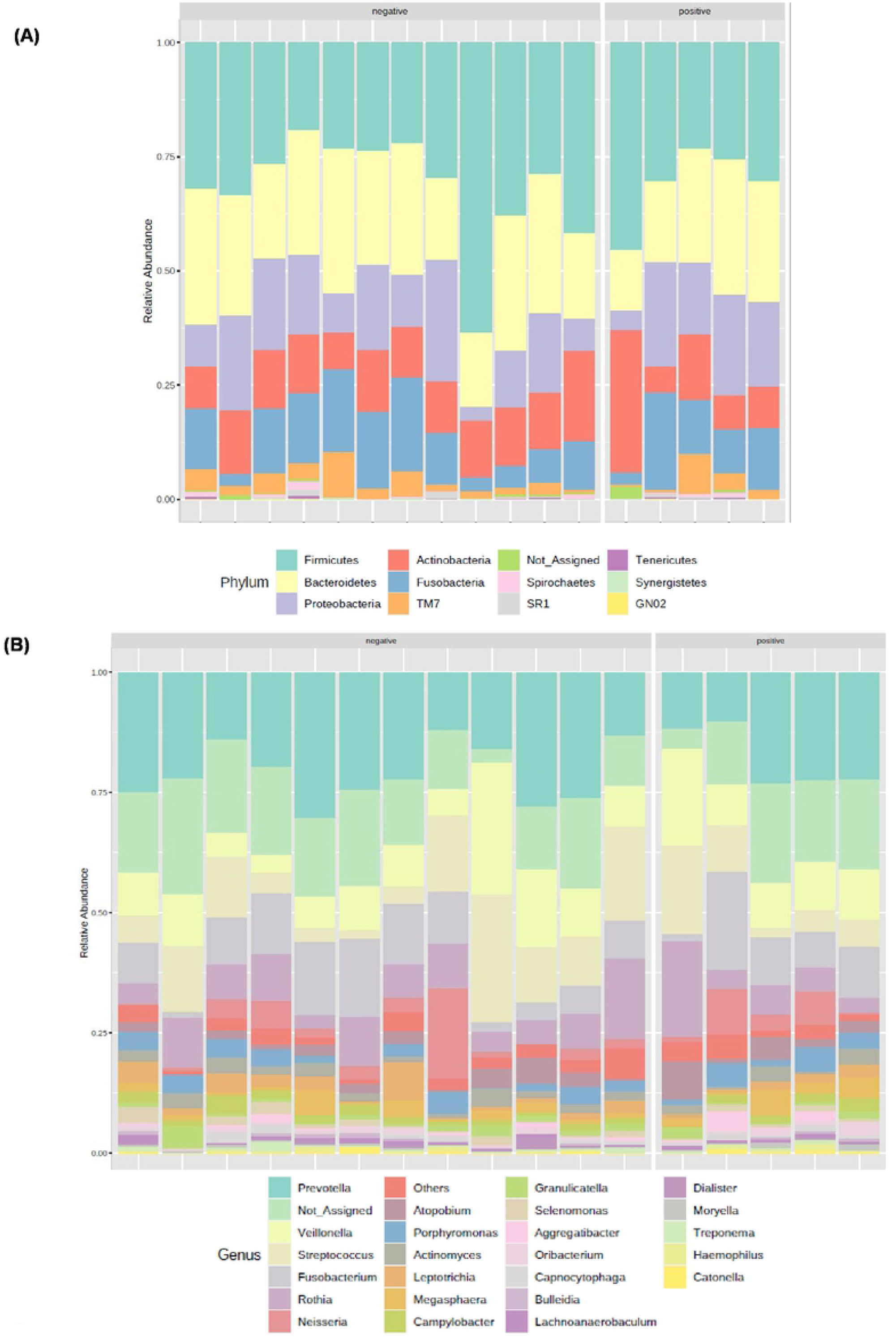
Bar plots of bacterial composition in the saliva samples. Samples grouped by HIV-negative (n=12) and positive (n=5). (A) at the phylum level. (B) at the genus level.

**S3 Fig.**
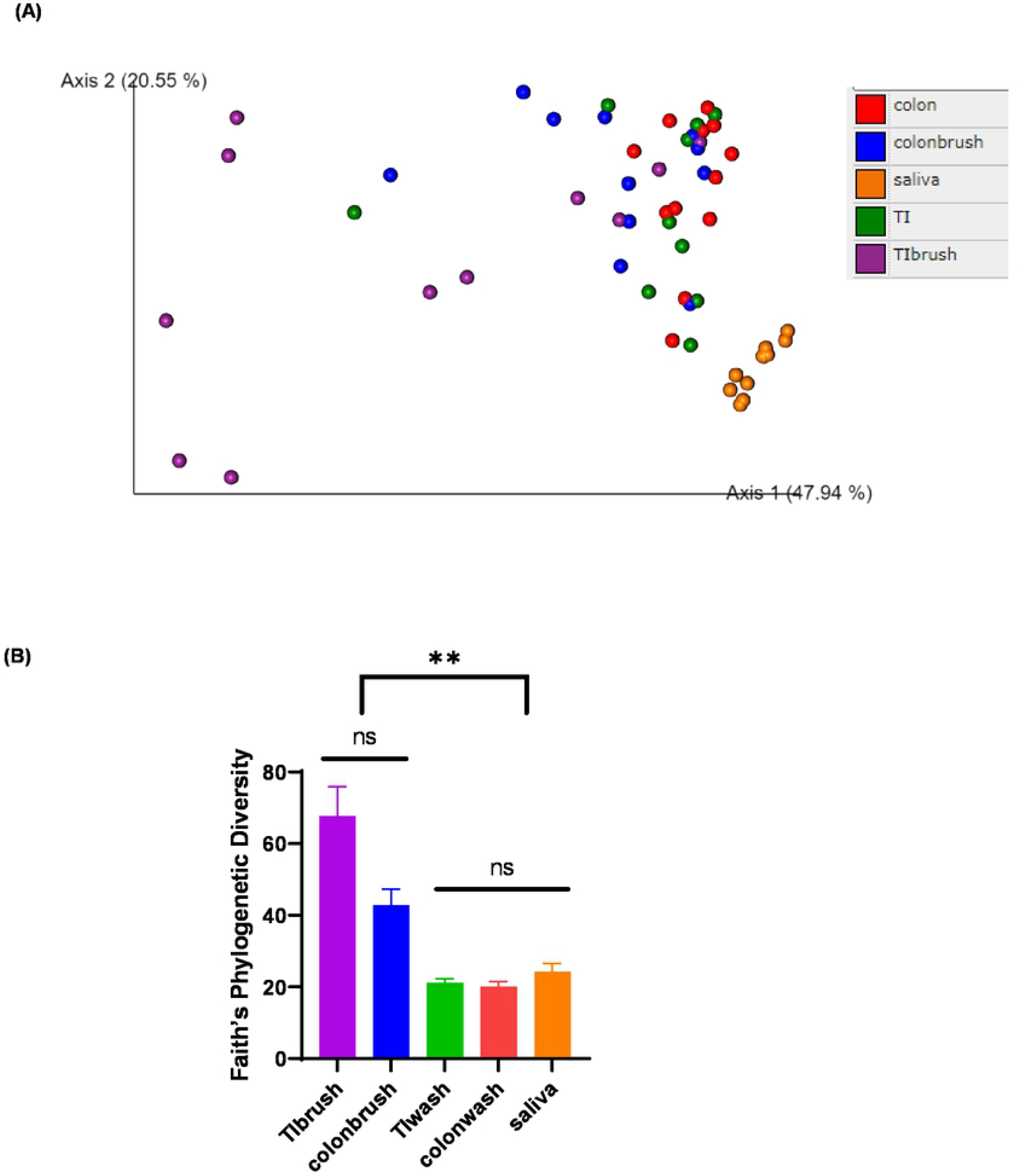
Diversity analysis of the HIV-negative samples. Principal coordinates analysis (PCoA) plot of weighted UniFrac distances (metrics of β -diversity). Samples grouped by TI wash (n=10), TI brush (n=11), colon wash (n=12), colon brush (n=11) and saliva (n=12). Falst discovery rate corrected q-value < 0.001 between saliva and every other group. q-value = 0.001 between TI brush and other intestinal samples (TI wash, colon wash, colon brush). q-value > 0.2 between colon, colon brush and TI wash. Faith Phylogenic Diversity (metrics of α-diversity) at sequencing depth 80000. Samples grouped by TI wash (n=10), TI brush (n=11), colon wash (n=12), colon brush (n=11) and saliva (n=12). “**” represents P< 0.001, “ns” represents not significant.

**S1 Table. Patients’ clinical characteristics**.

**S2 Table. Sequencing count of samples through each step in the DADA2 pipeline**.

**S3 Table. Sequencing count of amplicon sequence variants (ASVs) in all sample types**.

**S4 Table. Predicted pathway by PICRUSt**.

